# Confocal Live-Cell Imaging Under Ultralow Irradiance Using Superconducting Transition-Edge Sensor

**DOI:** 10.64898/2026.11.16.767203

**Authors:** Chigusa Okano, Chikaho Sano, Chihiro Horie, Yujiro Eto, Hiroaki Takaku, Kazuki Niwa, Daiji Fukuda, Nobuhiko Nomura, Yutaka Yawata

**Affiliations:** Institute of Life and Environmental Sciences, University of Tsukuba, Tsukuba, Ibaraki, Japan; Center for Science Adventure and Collaborative Research Advancement (SACRA), Graduate School of Science, Kyoto University, Kyoto, Kyoto, Japan; HIKARI-COOL Kyoto, Kyoto University Institute for Advanced Study, Kyoto, Kyoto, Japan; Graduate School of Life and Environmental Sciences, University of Tsukuba, Tsukuba, Ibaraki, Japan; National Institute of Advanced Industrial Science and Technology (AIST), Tsukuba, Ibaraki, Japan; Faculty of Applied Life Sciences, Niigata University of Pharmacy and Medical and Life Sciences, Niigata, Niigata, Japan; Microbiology Research Center for Sustainability (MiCS), Tsukuba Institute for Advanced Research (TIAR), University of Tsukuba, Tsukuba, Ibaraki, Japan

## Abstract

Confocal laser scanning fluorescence microscopy (CLSFM) is widely used for three-dimensional live-cell imaging. However, high irradiance is required to overcome dark counts in photodetectors, often leading to observation bias. In this study, we examine whether a CLSFM system equipped with a virtually dark-count-free superconducting transition-edge sensor (TES) can enable live-cell imaging under low irradiance, thereby mitigating observation bias. Using green fluorescent protein-tagged HeLa cells, we determined the minimum laser power required for live-cell imaging and demonstrated suppression of photobleaching and cell-division arrest during long-term observation. Innate fluorescence signature analysis revealed that TES-CLSFM could discriminate between two oleaginous yeast species with different lipid productivities at ca. 1/400 the irradiance required by conventional CLSFM. These results establish TES-CLSFM as a minimally invasive imaging modality for the quantitative physiological assessment of living cells.

## Introduction

Confocal laser scanning fluorescence microscopy (CLSFM) is one of the most widely used three-dimensional live-cell imaging modalities because it provides high axial resolution^1,2^. It has been applied to visualize diverse cellular responses to external stimuli, including thermal, electrical, and hormonal perturbations^3–5^. More recently, CLSFM has been extended to probe the physiology of intact cells through innate fluorescence (IF) signature analysis^6^. In this approach, the IF signature—an ensemble of weak autofluorescence emitted by intracellular molecules—is extracted in three dimensions at the single-cell level and used as an intrinsic biomarker for species identification, growth-phase assessment, and evaluation of metabolic productivity.

Achieving high-resolution CLSFM typically requires intense laser irradiation, both to obtain enough fluorescence photons from small focal volumes and to overcome detector dark counts—spurious detection events that occur even in the absence of light^7^. Conventional photodetectors widely used in CLSFM, such as photomultiplier tubes (PMTs), have dark counts of ca. 10–100 counts per second^[7,8]^. In PMTs, these dark counts arise primarily from thermionic emission. In conventional CLSFM, the irradiance at the focal plane is of the order of 10^6^–10^8^ W/m^2^, exceeding the solar constant by three to five orders of magnitude^1^. Such high irradiance can induce photobleaching and a various levels of non-lethal photodamage, which can manifest as observation bias—systematic deviations of cellular behavior from native responses during imaging. Reported examples include cell-cycle arrest, reduced migration of motor-neuron growth cones, calcium transients in chondrocytes, and loss of mitochondrial membrane potential^9,10^. Such observational bias can compromise long-term imaging and faithful physiological evaluation of living samples.

Here, we investigated whether a CLSFM system equipped with a virtually dark-count-free superconducting transition-edge sensor (TES-CLSFM) could enable long-term imaging and quantitative evaluation of living samples under low-irradiance conditions, thereby mitigating observation bias. We have recently proposed the concept of TES-CLSFM^11–13^. A TES is a microcalorimeter-type photodetector that resolves individual photon energies by measuring resistance changes at the superconducting transition edge^14,15^. Owing to its extremely low operating temperature, the TES exhibits near-zero dark counts (<1 Hz), making it well suited to photon-limited imaging. This capability allows the long-term accumulation of time-resolved photon signals and facilitates detection of faint fluorescence with sufficient quantitative reliability at reduced irradiance.

Using a genetically modified HeLa cell line stably expressing green fluorescent protein (GFP) we first determined the minimum laser power required for fluorescence-tag-assisted live-cell imaging using TES-CLSFM. We then examined how minimal irradiance affects photobleaching and cell-division arrest, a representative form of photodamage, during long-term imaging. Finally, we evaluated the utility of TES-CLSFM for the IF-based physiological assessment of intact cells under low irradiance by testing whether it could discriminate between two intact oleaginous yeasts with different lipid productivities.

The findings reveal that TES-CLSFM can visualize fluorescently tagged cells at irradiance levels comparable to the solar constant and suppress photobleaching and cell-division arrest during long-term imaging at approximately ten times the solar constant. TES-CLSFM-based IF analysis successfully classified the two oleaginous yeasts at ca. 1/400 the irradiance used in a previous study using conventional CLSFM. Collectively, these results indicate that TES-CLSFM provides a promising platform for minimally invasive physiological evaluation by effectively reducing the observation bias arising from photodamage.

Beyond the virtually dark-count-free characteristics focused on thus far, TES possesses the distinct advantage of a significantly broader detection bandwidth than conventional photodetectors, spanning the spectrum from visible light to near-infrared. Although our findings demonstrate its efficacy within the visible light spectrum—the most common spectral region for live-cell imaging—this broadband capability is expected to enable imaging in the near-infrared spectrum, which supports even less-invasive cell imaging.

## Results

### TES-CLSFM wavelength resolution and noise

The optical configuration of the TES-CLSFM system used here is presented in Fig. 1a. The system employs a fiber-coupled Ti/Au bilayer optical TES, optimized for ca. 1,550 nm. The optical fiber is connected to confocal optics, serving as both a pinhole and light carrier, sending fluorescence from the room-temperature optics to the TES within the adiabatic demagnetization refrigerator. To analyze the energy resolution of the TES-CLSFM, we measured the energy distribution of the pulsed laser and calculated the full width at half maximum (*E_FWHM_*) of the peak as the energy resolution of the TES (Fig. 1b), obtaining a resolution of 116 meV. Referring to that value, we estimated the wavelength resolution (*λ_FWHM_*) using Eq. 1 :

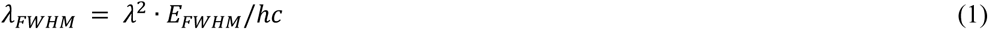

**Figure 1.**
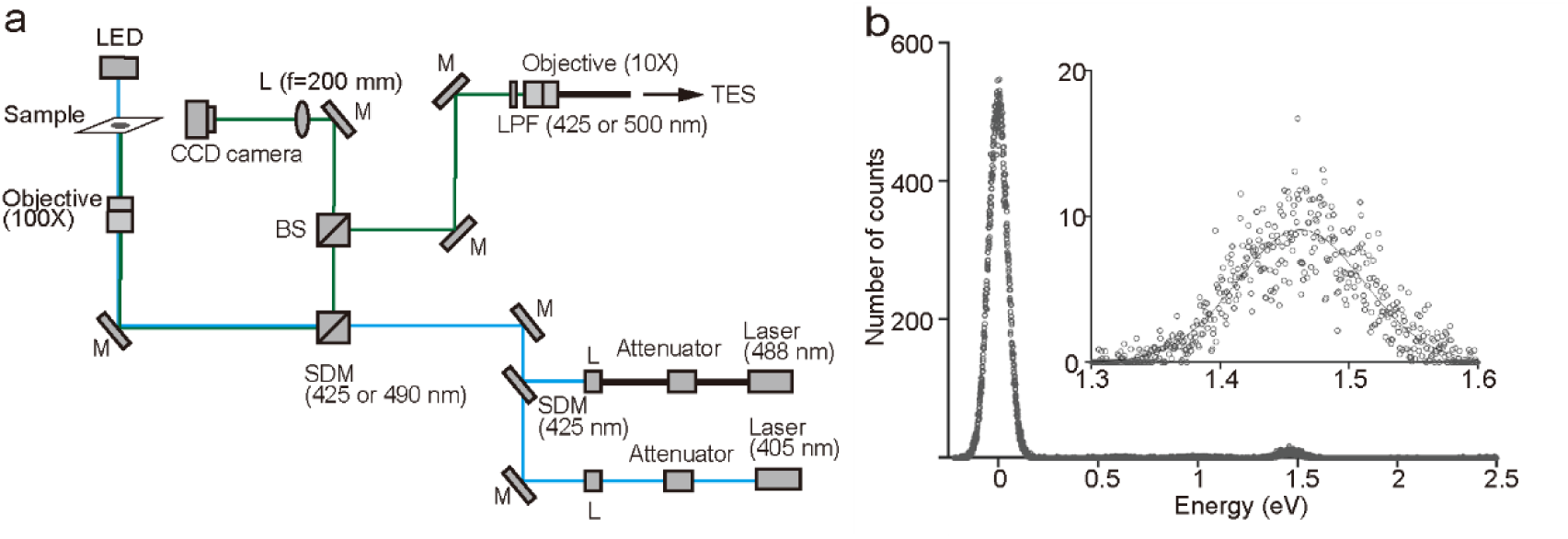
TES-CLSFM characteristics: Optical configuration and energy resolution. (a) Schematic diagram of the optical configuration in TES-CLSFM. M: Mirror; L: Lens; BS: Beam splitter; SDM: Shortpass dichroic mirror; LPF: Longpass filter. Blue and green lines, excitation and fluorescence optical paths, respectively. (b) Energy distribution of a pulsed laser at 848 nm (a central wavelength), measured using the TES. The TES was directly irradiated by laser at 2 kHz through an optical fiber. Horizontal and vertical axes, photon energy and detection count, respectively. Total count; 10,000. The detection peak at 0 eV corresponds to zero photons, while that at 1.46 eV corresponds to one photon (inset: enlarged view of this peak). Solid line, fitted Gaussian distribution.

where *λ* is wavelength of incident light, *h* is Planck’s constant, and *c* is the speed of light^12^.

For a wavelength of 550 nm—the middle wavelength in the visible light region—*λ_FWHM_* was 28 nm. This indicates that TES-CLSFM provides spectroscopy with a wavelength resolution ca. 1/20 that of the wavelength in the visible light region.

Next, we quantified the intrinsic noise level of the TES. The pulse-height distribution of dark events was acquired under light-tight conditions, implemented by disconnecting the optical fiber and optically sealing the TES fiber port. Over 50,000 s, 19 events were recorded above the zero-photon state, corresponding to a dark count rate of 3.8×10^−4^ counts s^−1^. This ultralow rate is consistent with previous reports that TES detectors exhibit dark counts below 1 Hz, more than four orders of magnitude lower than those of typical PMTs^7,8^. The confirms that the TES operates in a virtually dark-count-free regime under the conditions used here, motivating the subsequent determination of the minimum excitation power required to achieve a sufficient signal-to-noise ratio (SNR) for live-cell imaging via TES-CLSFM.

### Minimum TES-CLSFM laser power for fluorescent-tagging assisted live-cell imaging

We examined the order of magnitude of the minimum laser power required for cell visualization and quantitative fluorescence analysis via TES-CLSFM. Genetically modified HeLa cells stably expressing GFP (the HeLa/GFP cell line) were placed on the TES-CLSFM device, and a 50 × 50 µm area was scanned at 50 ms per pixel while varying the laser power from 0.33 to 35 nW. For visualization, we define the minimum laser power as the lowest level at which the SNR reached or exceeded 10. The TES-CLSFM successfully visualized the cells even at 0.33 nW (Fig. 2a), with an SNR of 10.02 at this power, thus identifying 0.33 nW as the minimum laser power required for visualization (Fig. 2b). To determine the minimum laser power required for quantitative fluorescence analysis, we assessed whether the histogram of fluorescent photons for a single cell followed a Gaussian distribution. For an irradiation power as low as 0.33 nW, the histogram was already well fitted to Gaussian distribution, except within the lower-most tail, corresponding to less than −1.72 σ (i.e., 4.3% of the area under the curve). The histogram for irradiation at ≥3.5 nW also captured the entire distribution, suggesting that 3.5 nW is sufficient for quantitative fluorescence analysis. These results indicate that 0.33 nW is sufficient for visualization, whereas 3.5 nW suffices for quantitative fluorescence analysis using TES-CLSFM. Based on these findings, compared with conventional confocal microscopy measurements performed at laser powers of 5–10 µW^16^, TES-CLSFM enables both observation and quantitative analysis at laser powers that are orders of magnitude lower.

**Figure 2.**
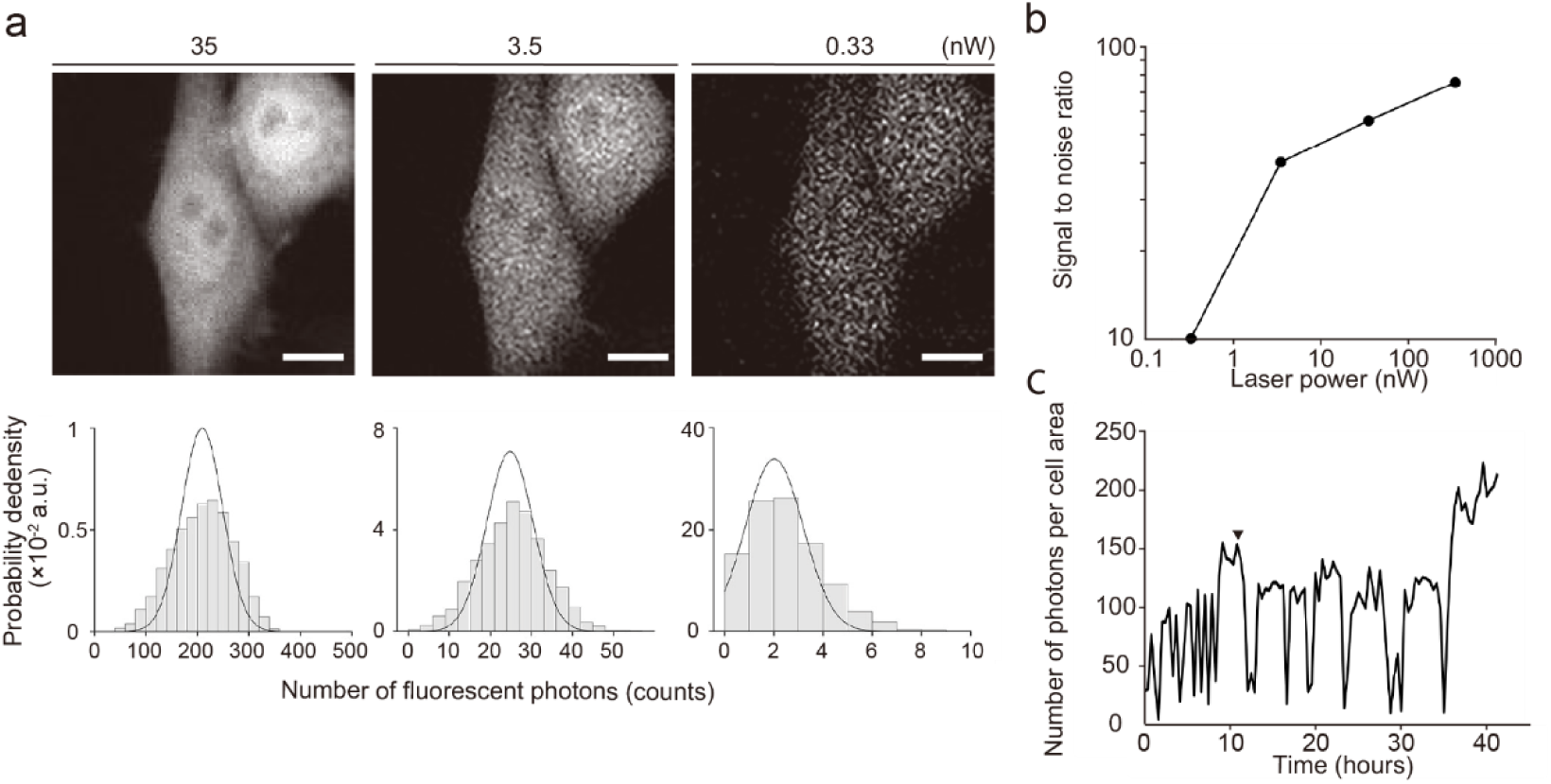
Exploration of minimal irradiation dose and long-term fluorescence observation of HeLa/GFP cells. (a) Upper panel: Optical slice images of HeLa/GFP cells (Scale bar: 10 µm). Lower panel: Histogram of fluorescence intensity within a cell region (corresponding to left cell in upper panel). Example data; similar trends were observed in at least five cells. (b) Signal-to-noise ratio as a function of laser power. (c) Temporal dynamics of fluorescence intensity within a representative cell. Irradiation dose, 3.5 nW; duration, 41 h; time interval, 16.6 min. Triangle, timepoint of cell division. Throughout the figure: Excitation laser wavelength, 488 nm; detection wavelength, 500–650 nm; exposure time per pixel, 50 ms. Lens magnification, 100×.

### Evaluation of observation bias under long-term observation

To evaluate the feasibility of long-term time-lapse imaging using TES-CLSFM, we performed time-lapse imaging of HeLa/GFP cells for 41 h and assessed potential observation bias in terms of cell proliferation and fluorescence photobleaching. Cells were held in a stage incubator, and 2D scanning images were taken at intervals of 16.5 min via laser irradiation at 3.5 nW. In a prior study^17^, HeLa cells exhibited cell-division arrest under specific light-irradiation conditions (excitation, 514 nm; 2.5×10^7^ W/m^2^; for 240 s). We confirmed that continuous two-dimensional imaging of HeLa/GFP cells using a conventional confocal microscope (excitation: 405 nm; 9.6×10^7^ W/m^2^) induced both inhibition of cell proliferation (Supplementary Video 1) and an irreversible reduction in fluorescence intensity (Supplementary Fig. 1). Although the excitation wavelength, irradiation interval, and irradiation intensity differ from those used in TES-CLSFM, these results indicate that HeLa/GFP cells are susceptible to phototoxicity-induced proliferation inhibition and irreversible photobleaching under conventional imaging conditions at a comparable SNR (109 ± 19.4, n = 4). In contrast, our time series images confirmed continuous cell proliferation (Supplementary Video 2). This suggests that the irradiation conditions used in this study did not induce photodamage severe enough to inhibit cell division.

Next, we tracked the temporal dynamics of the average fluorescence intensity for a representative single cell via image analysis (Fig. 2c). In our system, although fluorescence intensity fluctuated over time, a monotonic decline in emission intensity, indicative of irreversible photobleaching, was not observed. The observed fluctuations in fluorescence intensity can be explained by two factors. The first reflects morphological dynamics along the Z-axis. When HeLa/GFP cells adhere to a glass surface, they exhibit pronounced morphological fluctuations associated with cytoskeletal dynamics and cell cycle–dependent structural changes. The second reflects fluctuations in *gfp* expression in accordance with the cell cycle. In the HeLa/GFP cells used here, *gfp* expression is driven by a cytomegalovirus promoter, which has been reported to exhibit higher transcriptional activity during the S phase than during the G0/G1 phase^18^. Both photodamage and photobleaching can involve reactive oxygen species (ROS)-mediated oxidative reactions affecting cellular components and fluorescent chromophores, respectively^9,19^. Therefore, the minimal levels of photodamage and photobleaching observed here indicate that the irradiation settings applied here effectively suppressed ROS generation. Together, these observations indicate that TES-CLSFM enables extended time-lapse live-cell imaging with minimal ROS-mediated observation bias.

### Physiological evaluation of intact cells via TES-CLSFM under nanowatt-order irradiation

Finally, we examined whether TES-CLSFM allows the physiological evaluation (via single-cell autofluorescence analysis) of intact cells under low-power laser irradiation. We acquired IF images of oleaginous yeast strains using 119 nW laser irradiation (excitation wavelength: 405 nm; exposure time: 50 ms/pixel), trained a machine-learning model to classify the two strains with different lipid production capacities, and evaluated their performance. TES-CLSFM successfully captured IF images of these strains at 119 nW laser irradiation (Fig. 3a). Using image analysis, the contours of individual cells were identified and a set of eight emission bands were extracted from each cell region as IF signatures (Fig. 3b). We used two machine learning models to classify the IF signatures: logistic regression and linear discriminant analysis (LDA), both using the first two principal components (PC1 and PC2) obtained from principal component analysis (PCA) as input features. In the two-dimensional PCA plot of all 90 samples, the two yeast strains formed distinct clusters, with the decision boundary (determined using logistic regression) effectively separating the clusters (Fig. 3c). Five-fold cross-validation yielded classification accuracies of 93.3 ± 4.6% and 91.1 ± 3.0% for logistic regression and LDA, respectively. This accuracy is comparable to the previously reported accuracy of 80.1–85.7% for three-class discrimination using LDA^20^, in which IF images were obtained using conventional CLSFM with four excitation wavelengths, including 405 nm excitation at a laser power of 38 µW. The receiver operating characteristic (ROC) curves^21^ obtained through five-fold cross-validation for the logistic regression and LDA-based classification (Extended Data Figs 1 and 2) exhibit areas under the curve (AUC)^22^ of 0.980 and 0.950, respectively, with standard deviations of 0.038 and 0.050. These high AUC scores demonstrate that both models achieved reliable classification. These results suggest that TES-CLSFM is well suited for minimally invasive and tag-free physiological classification.

**Figure 3.**
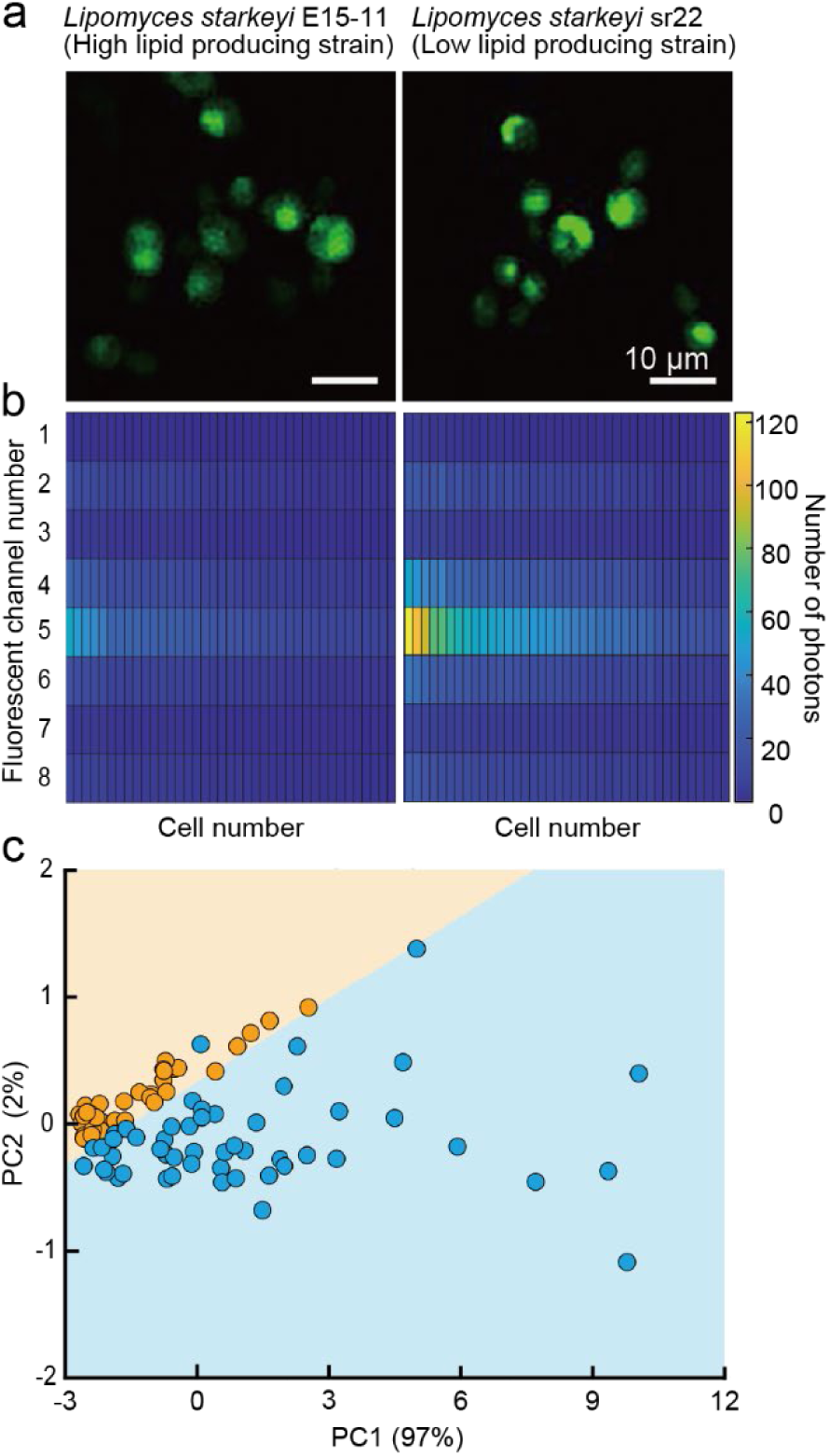
Single-cell innate fluorescence signatures of two oleaginous yeast strains using TES-CLSFM. (a) Optical slice images of *Lipomyces starkeyi* E15-11 and sr22 at representative fluorescence channels (580–630 nm), in pseudo color. E15-11 and sr22 are high lipid producing and low lipid producing strains, respectively^28,29^. Laser wavelength, 405 nm; irradiation dose, 119 nW; lens magnification, 100×; exposure time per pixel, 50 ms. (b) Heatmap of 39 representative single-cell IF signatures. Each column denotes one cell, and each row a fluorescence channel. Channels: 1, <430 nm; 2, 430–480 nm; 3, 480–530 nm; 4, 530–580 nm; 5, 580–630 nm; 6, 630–680 nm; 7, 680–730 nm; and 8, 730–780 nm. Cell color, number of detected fluorescent photons. Fluorescence intensity was not corrected for TES wavelength sensitivity. (c) Principal component analysis projection of single-cell IF signatures in the two strains. Orange and blue points, strains E15-11 (n=39) and sr22 (n=51), respectively. Horizontal and vertical axes, first principal component (PC1) and second principal component (PC2), respectively. Boundary between the blue and orange regions, decision boundary, determined using logistic regression.

## Discussion

Our findings demonstrate the noise-free characteristics of the TES and reveal that TES-CLSFM exhibits a spectral resolution of ca. 1/20 that of the wavelengths in the visible light region. Based on live-cell imaging under varying irradiance levels, TES-CLSFM is capable of visualizing and quantitatively evaluating cells at much lower laser power than conventional CLSFM. Long-term imaging analysis revealed that TES-CLSFM induced minimal photodamage. TES-CLSFM enabled IF-based classification with a high accuracy comparable to that of conventional CLSFM, even under nanowatt laser power conditions.

Conventional diffraction-grating-based CLSFM commonly achieves a spectral resolution of 2.5 nm (Nikon Corporation. A1HD25/A1RHD25 Confocal Microscope Specifications. Nikon Healthcare. https://www.microscope.healthcare.nikon.com/en_EU/products/confocal-microscopes/a1hd25-a1rhd25/specifications), whereas the TES-CLSFM exhibited a spectral resolution of 28 nm at 550 nm. Therefore, the spectral resolution of the TES-CLSFM is 11 times lower than that of the conventional CLSFM at 550 nm. Although our IF analysis revealed that this spectral resolution was sufficient for IF classification, the relatively low spectral resolution may preclude the use of TES-CLSFM in applications requiring high-spectral resolution, such as Raman microscopy. Although lowering the critical temperature can improve the spectral resolution, it can also reduce the response speed²³, thereby reducing the overall scanning rate.

Here, the TES-CLSFM successfully imaged living HeLa/GFP cells even at 0.33 nW. Irradiance (*I*) is defined as the laser power divided by the area of the focal spot, as in Eq. 2:

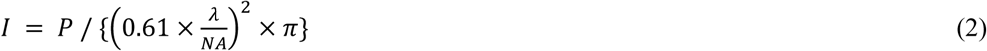

where *P*, *λ*, and *NA* are the laser power (unit: Watt), laser wavelength, and the numerical aperture of objective lens, respectively. The estimated irradiance for 0.33 nW was 0.96 × 10^3^ W/m^2^ (*λ*, 488 nm; *NA*, 0.9; Lens: PlanN 100×; Olympus, Tokyo, Japan), suggesting that the minimum irradiance for fluorescence-tagging imaging is comparable to the solar constant^1^ (1.36 × 10^3^ W/m^2^). The estimated irradiance of conventional CLSFM imaging of HeLa/GFP cells^16^ is 5.24 × 10^7^ W/m^2^.

TES-CLSFM has the potential to achieve even lower irradiance. Because the current optical TES is optimized for the near-infrared region (at ca. 1,550 nm)^[11]^, its sensitivity in the visible light region is <0.1. If the optical cavity of the TES is optimized for the visible light region and for the quantum efficiency of the TES, the irradiance required in TES-CLSFM will decline even further, while maintaining the same image quality. Considering that the maximum detection efficiency of a TES with an optical cavity has been reported^15^ to be 98%, optimizing the optical cavity is expected to be highly effective for imaging under even lower irradiances.

An extremely long exposure time (50 ms/pixel) allows the TES-CLSFM to achieve live-cell imaging under extremely low irradiance. In microscopy, in order to capture the full distribution of fluorescence intensity under low irradiance, a long exposure time is typically required to sufficiently accumulate temporally discrete fluorescence signals. While common single-photon detectors such as PMTs undesirably mask discrete signals with their high dark counts, the virtually dark-count-free TES-CLSFM system can extend the exposure time almost indefinitely and allows further reductions irradiance.

For TES-CLSFM, the total scan time increased linearly with the exposure time, thus reducing temporal resolution. In conventional CLSFM, the exposure time ranges from one to tens of microseconds per pixel (https://www.microscope.healthcare.nikon.com/en_EU/products/confocal-microscopes/a1hd25-a1rhd25/specifications), whereas the exposure time achieved here was 50 ms per pixel, resulting in an exceptionally long scan time of ca. 8.3 min for a 100 × 100 pixel field of view. To mitigate this long exposure time, the detection efficiency at the target wavelength could be optimized or a TES array could be used. As already discussed, optimizing TES detection efficiency for the target fluorescence wavelength range reduces the required exposure time, while preserving image quality under the same irradiance. Optical TES arrays provide a promising tool for simultaneously achieving a long exposure time per pixel and short total scan time^24^. An array of TES devices can be integrated on a single chip. By combining a TES array with multipoint CLSFM, the total scan time can be reduced linearly in proportion to the number of TES detectors in the array.

The total photon energy (in J) received by the sample during TES-CLSFM imaging is not necessarily lower than that received under conventional CLSFM, because the increased exposure time offsets the low irradiance. Nevertheless, from the perspective of phototoxicity, it is noteworthy that TES-CLSFM enables live-cell imaging at irradiance levels ca. four orders of magnitude lower than those allowed by conventional CLSFM. Under conditions of equal total energy, it is well-established that lower irradiance combined with longer exposure times causes less biological photodamage^9^. Consistent with this, our long-term imaging under low irradiance did not lead to an irreversible decline in fluorescence intensity (irreversible photobleaching). This intrinsic capacity to avoid photobleaching highlights the advantage of TES-CLSFM for non-invasive physiological evaluation and may enable tracking of intracellular molecular dynamics while maintaining low expression levels of fluorescent molecules.

Here, TES-CLSFM using laser irradiation of 119 nW enabled IF-based physiological evaluation of intact cells, with an estimated irradiance of 5.0 × 10^5^ W/m^2^, ca. 1/400 that estimated for a previous study in which the same oleaginous yeast strains were imaged using conventional CLSFM^20^ (assuming an objective NA of 0.95). The same study also revealed that *L*. *starkeyi* wild-type cells are in the lipid-accumulation phase during days 1–3 of cultivation, and that lipid accumulation on day 1 is not readily detectable by microscopy. This suggests that IF on day 1 of cultivation is more likely to reflect differences in metabolic state rather than differences in lipid content. Here, machine learning-based classification of the yeasts revealed that TES-CLSFM supports high-accuracy classification, equivalent to conventional CLSFM, under low irradiance. TES-CLSFM-based physiological evaluation may therefore enable minimally invasive, culture-free, and real-time screening of specific cells from intact cell populations.

In recent years, 3D time-lapse imaging combined with fluorescence-tagging technology has successfully revealed the spatiotemporal dynamics of cell lineages in multicellular structures such as embryos and bacterial biofilm^25,26^. However, such studies suffer from low spatial resolution, owing to the use of light-sheet fluorescence microscopy to maintain low irradiation power. As TES-CLSFM can overcome high irradiance, it has the potential to achieve high spatial resolution, thus supporting the analysis of spatiotemporal dynamics. Furthermore, in combination with single-cell IF signature analysis, TES-CLSFM may support the tag-free identification of cell lineages not only for microorganisms, but also for a wide range of organisms such as plants and mammals, because autofluorescence is present across biological species^27^. TES-CLSFM therefore provides a powerful tool for exploring the spatiotemporal dynamics of cell lineages in a minimally invasive manner and with minimal observation bias. This is particularly appropriate for the delicate samples used in regenerative medicine, such as iPS and ES cells, which are highly susceptible to phototoxicity.

## Funding

This work was supported in part by grants from JST CREST to D. Fukuda (grant number JPMJCR17N4), JST ERATO to N. Nomura (grant number JPMJER1502), MEXT Leading Initiative Excellent Young Researchers to Y. Yawata, JST FOREST to Y. Yawata (grant number JPMJFR210G), JSPS, JSPS Research Fellowships for Young Scientists to C. Okano (grant number 23KJ0289), and from the RIKEN TRIP initiative.

## Acknowledgements

We thank Kyosuke Takabe for technical assistance with image processing using MATLAB. We thank Ayumi Hiroki, Azumi Kumagai, and Shiomi Yawata for technical support with cell culture. We thank Takahiro Kikuchi and Koki Shirota for helpful discussions.

## Author contributions

Y.Y., D.F., N.N., and C.O. designed the experiments. D.F. constructed the TES device. C.O., D.F., and Y.E. designed and constructed the CLSFM-TES optical system. K.N. constructed the data-acquiring system. H.T. contributed the yeast strains and cultures. C.O., D.F., C.S., and C.H. performed the observations and experiments. C.O. analyzed the data. C.O. and Y.Y. drafted the manuscript. All of the authors have reviewed the manuscript.

## Competing interests

The authors declare no competing interests.

## Data availability

The data are available from the author upon reasonable request.

## Methods

### Fabrication of TES device

The TES device was constructed as previously described^12^. A bilayer Ti/Au superconducting film was incorporated into an optical cavity comprising multilayer dielectric films. The optical cavity was designed and optimized for a wavelength of 1,550 nm. The TES device was mounted on the cold stage of an adiabatic demagnetization refrigerator at 100 mK. Fluorescent photons from the microscope are delivered to the TES via an optical fiber (780HP; Thorlabs, Newton, NJ, USA; mode field diameter: 5.0 μm at 850 nm) connected to the optical cavity. The photon energy is absorbed by the TES, causing electrical resistance to increase. This change in resistance is converted into a voltage signal using a superconducting quantum interference device amplifier placed near the TES. The pulse height distribution of the signal was analyzed using either an oscilloscope (MSO64; Tektronix, Beaverton, OR, USA) or a multichannel analyzer (MCA7600; SEIKO EG&G, Tokyo, Japan). For imaging measurements, a single-photon counter (SR400; Stanford Research Systems, Sunnyvale, CA, USA) was used to count pulses within a predefined pulse height range.

### Construction of TES-CLSFM

The microscope optical system and the adiabatic demagnetization refrigerator, in which the TES device was mounted, were placed separately and connected via an optical fiber. The microscope optics were assembled inside a light-tight enclosure to maintain the dark environment. Within the microscope, most of the light propagates through free space. Two excitation light paths, optimized at 405 nm and 488 nm, respectively, were manually switched by inserting a mirror. The beams from the excitation lasers (LP405-SF10 and LP488-SF20; Thorlabs, Newton, NJ, USA) are collimated using a lens (PAF2-A7A or PAF2P-A15A; Thorlabs, Newton, NJ, USA) and introduced into the optical path via a dichroic or standard mirror. The beams are then directed at an objective lens (PlanN 100×; Olympus, Tokyo, Japan; numerical aperture, 0.9) through a dichroic mirror (DMSP425 or DMSP490; Thorlabs, Newton, NJ, USA) and focused on the sample. A stage-top CO_2_ incubator (Tokai Hit, Shizuoka, Japan) was installed on a custom aluminum stage mounted on an electronically controlled XYZ actuator to enable automated XY scanning and precise focusing. A LED provides transmitted illumination for sample alignment. A beam splitter (BS028; Thorlabs, Newton, NJ, USA), located between the dichroic mirror and long-pass filter, directs the transmitted light from the sample to a CCD camera (DMK33UX174; The Imaging Source, Bremen, Germany). The fluorescent photons emitted by the sample pass through the dichroic mirror, a longpass filter with an optical density of 4 (Edmund Optics, Barrington, NJ, USA), and a second objective lens (M Plan Apo 10×; Olympus, Tokyo, Japan) and are coupled into the optical fiber connected to the TES. This fiber serves as both a pinhole for confocal microscopy and a light guide to deliver the photons to the TES. The entire system is controlled using LabVIEW (National Instruments, Austin, TX, USA).

### TES-CLSFM characterization

The energy distribution of the 848 nm laser pulses was acquired by directly irradiating the TES with the laser through the optical fiber, and the energy resolution was determined as the full width at half maximum (*E_FWHM_*) of the peak corresponding to the single-photon state; *E_FWHM_* was calculated by fitting the peak to a Gaussian model using the *fit* function in MATLAB (MathWorks, Natick, MA, USA). To examine the dark counts of the TES-CLSFM system, the energy distribution was measured over 50,000 s with the fiber port sealed.

### Cell culture conditions

The HeLa/GFP cell line (Cell Biolabs, San Diego, CA, USA) and two *Lipomyces starkeyi* strains, E15-11 (a high lipid producing strain) and sr22 (low lipid producing strain) were used^28,29^. The HeLa/GFP cells were cultured in D-MEM (without L-glutamine or phenol red; Fujifilm Wako Pure Chemical, Osaka, Japan) supplemented with 9.7% (v/v) fetal bovine serum (Thermo Fisher Scientific, Waltham, MA, USA), 1.9 mM L-glutamine (Fujifilm Wako Pure Chemical, Osaka, Japan), 1% (v/v) penicillin– streptomycin solution (100×; Fujifilm Wako Pure Chemical, Osaka, Japan), and 1% (v/v) MEM non-essential amino acids solution (100×; Merck, Darmstadt, Germany) on a glass bottom dish (AGC Techno Glass, Shizuoka, Japan) at 5% CO_2_ and 37 °C for 1 d. For microscopy, the cells were washed twice with fresh medium. During confocal imaging, the cells were maintained at 37 °C and 5% CO_2_ using the aforementioned stage incubator.

The oleaginous yeast strains were cultured as previously reported^20^. Briefly, the two strains were cultured on YPD agar plates at 30 °C for 4 d. The cells were collected from the YPD agar plates and inoculated into semi-defined glycerol liquid medium [0.5% (NH_4_)_2_SO_4_ (Fujifilm Wako Pure Chemical, Osaka, Japan), 0.1% KH_2_PO_4_ (Fujifilm Wako Pure Chemical, Osaka, Japan), 0.01% NaCl (Fujifilm Wako Pure Chemical, Osaka, Japan), 0.1% yeast extract (Kyokuto, Tokyo, Japan), 2% glycerol (Fujifilm Wako Pure Chemical, Osaka, Japan), 0.05% MgSO_4_·7H_2_O (Fujifilm Wako Pure Chemical, Osaka, Japan), and 0.01% CaCl_2_·2H_2_O (Fujifilm Wako Pure Chemical, Osaka, Japan)] in baffled flasks, and cultured at 30 °C for 72 h with shaking at 160 rpm. Next, these cells were inoculated into semi-defined medium [0.5% (NH_4_)_2_SO_4_ (Fujifilm Wako Pure Chemical, Osaka, Japan), 0.1% KH_2_PO_4_ (Fujifilm Wako Pure Chemical, Osaka, Japan), 0.01% NaCl (Fujifilm Wako Pure Chemical, Osaka, Japan), 0.1% yeast extract (Kyokuto, Tokyo, Japan), 5% glucose (Fujifilm Wako Pure Chemical, Osaka, Japan), 0.05% MgSO_4_·7H_2_O (Fujifilm Wako Pure Chemical, Osaka, Japan), and 0.01% CaCl_2_·2H_2_O (Fujifilm Wako Pure Chemical, Osaka, Japan)] in baffled flasks at 3.0×10⁶ cells/mL, and cultured at 30 °C for 24 h with shaking at 120 rpm. Finally, these cells were inoculated into semi-defined medium in baffled flasks at a density of 1.2×10⁷ cells/mL and cultured at 30 °C with shaking at 120 rpm. For observation, cells from the final culture were washed twice with PBS on the day of seeding and placed between an agar slab and the coverslip of a glass-bottom dish to maintain their position under moist conditions.

### TES-CLSFM microscopy of Live Cells

Coarse focusing was performed using bright-field imaging with epi-illumination using an LED lamp. For precise focusing, we searched for the optimal height (Z-axis position) at which the fluorescent signals were maximized, while checking the number of photons detected in the TES via confocal observation. The samples were then mechanically scanned along the X and Y axes using a motorized stage, with a pixel size of 0.5 μm and an exposure time of 50 ms per pixel. The HeLa/GFP cells and oleaginous yeasts were irradiated using 488 nm and 405 nm lasers, respectively. The detected emission wavelength range for HeLa/GFP cells was 500–650 nm. For yeasts, the emission wavelengths within the range of 430–780 nm were divided into eight channels with a 50 nm bandwidth. The irradiation power at the focal plane was measured using a power meter (PM-400; Thorlabs, Newton, NJ, USA) equipped with a slide-type sensor (S170C; Thorlabs, Newton, NJ, USA) and was adjusted using an electronic variable optical attenuator (V450F; Thorlabs, Newton, NJ, USA) or neutral density filter (Sigma Koki, Tokyo, Japan).

### Image analysis for HeLa/GFP cells

For the HeLa/GFP cells, the number of photons in the cell region was analyzed using a custom MATLAB routine. The cell region was manually defined and segmented into cell masks. The SNR was calculated as the difference between the mean intensity within the cell region and the mean background intensity, divided by the standard deviation of the background intensity. The distribution of the number of photons in the cell mask was visualized as a probability histogram using the ‘histogram’ function in MATLAB. The histogram was fitted to a Gaussian distribution using the ‘fitdist’ function in MATLAB.

For visualization, the display contrast of fluorescence images and videos was adjusted in Fiji (ImageJ) using identical minimum and maximum intensity settings across the images being compared. All quantitative analyses were performed using the original, unprocessed data.

### Reconstruction and analysis of single-cell IF signature

The single-cell IF signature was reconstructed using a custom MATLAB routine, with slight modifications from the described method^6^. Cell masks were created as described. The number of photons detected in each channel was linked to each cell mask, averaged over the mask, and organized into an eight-element matrix representing excitation–emission combinations, thus providing a single-cell IF signature.

Classification analysis was performed using a custom MATLAB routine. For visualization, all features were standardized to zero mean and unit variance before analysis. PCA was applied for dimensionality reduction using the ‘pca’ function, and the first two principal components (PC1, PC2) were used to visualize the distribution of the samples. Logistic regression models were fitted using the ‘mnrfit’ function, and decision boundaries were visualized using two-dimensional PCA plots. Classification performance was evaluated using stratified five-fold cross-validation. Within each fold, feature standardization and PCA were performed using the training data only. The test data were standardized using the mean and standard deviation derived from the corresponding training set and subsequently projected onto the PCA space derived from the training data. PC1 and PC2 were then used as input features for logistic regression and LDA. LDA was performed using the ‘fitcdiscr’ function. Classification accuracy was calculated for each test fold. The receiver operating characteristic (ROC) curves and areas under the curve (AUC) were computed using the ‘perfcurve’ function. Mean classification accuracies and AUCs, together with their standard deviations, were calculated across the five folds.

For visualization, the display contrast of fluorescence images and videos was adjusted in Fiji (ImageJ) using identical minimum and maximum intensity settings across the images being compared. All quantitative analyses were performed using the original, unprocessed data.

## Extended data figures

**Extended Data Figure 1.**
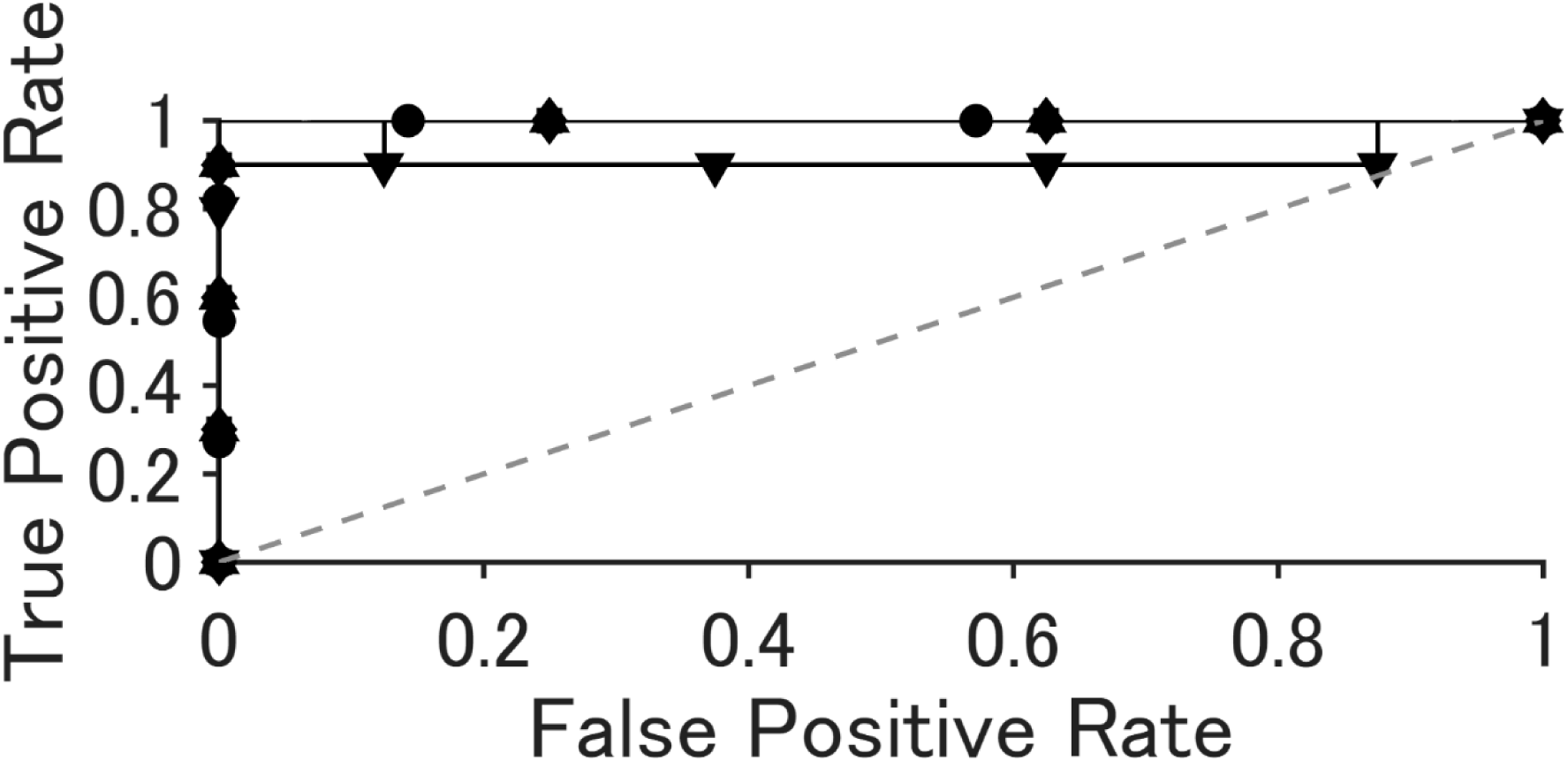
ROC curves obtained via five-fold cross-validation of logistic regression following PCA. Each curve represents an individual cross-validation fold. The dashed diagonal line indicates the performance expected from random classification.

**Extended Data Figure 2.**
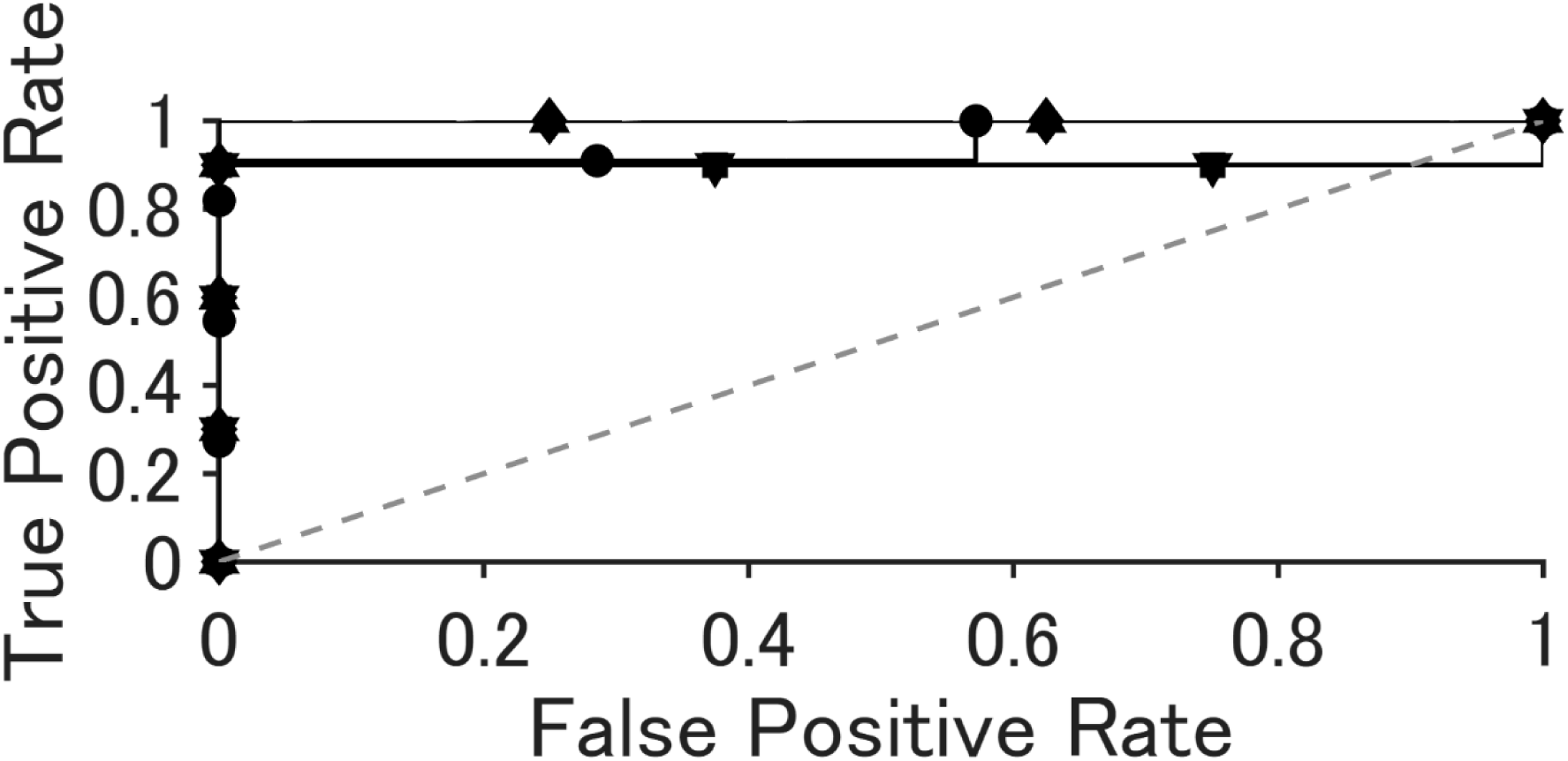
ROC curves obtained via five-fold cross-validation of LDA following PCA. Each curve represents an individual cross-validation fold. The dashed diagonal line indicates the performance expected from random classification.

## Notes

### Competing Interest Statement

The authors have declared no competing interest.

