## Supplementary Information for "Confocal Live-Cell Imaging Under Ultralow Irradiance Using Superconducting Transition-Edge Sensor"

### Supplementary Figures

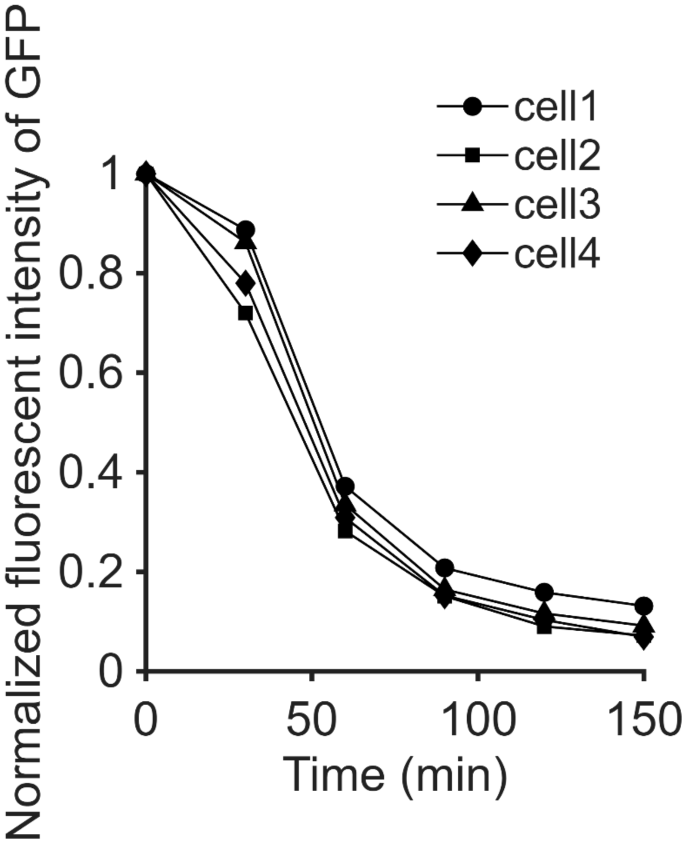


Supplementary Figure1

Temporal changes in fluorescence intensity of HeLa/GFP cells during continuous imaging with a conventional confocal microscope. Images were acquired using a 100× objective at an excitation wavelength of 405 nm and an excitation power of 8.75 µW, with an image size of 1024 × 1024 pixels and an exposure time of 28.85 µs per pixel. Continuous illumination was applied to the same field of view without an inter-frame interval.

### Supplementary Videos

Supplementary Video1

Long-term confocal imaging of HeLa/GFP cells using a conventional confocal microscope under continuous illumination. Images were acquired using a 100× objective at 405 nm excitation (8.75 µW), with an image size of 1024 × 1024 pixels and an exposure time of 28.85 µs per pixel. Imaging was performed in the same field of view without an inter-frame interval. The video was generated by extracting images acquired at 0, 30, 60, 90, 120, and 150 min after the start of imaging. (Related to Supplementary Fig. 1.)

Supplementary Video2

Long-term confocal imaging of HeLa/GFP cells using the TES-CFLSM.
